## Supplementary figures and images for "Non-thalamic origin of zebrafish sensory relay nucleus: convergent evolution of visual pathways in amniotes and teleosts"

### Supplementary file 1

#### Supplementary file 1

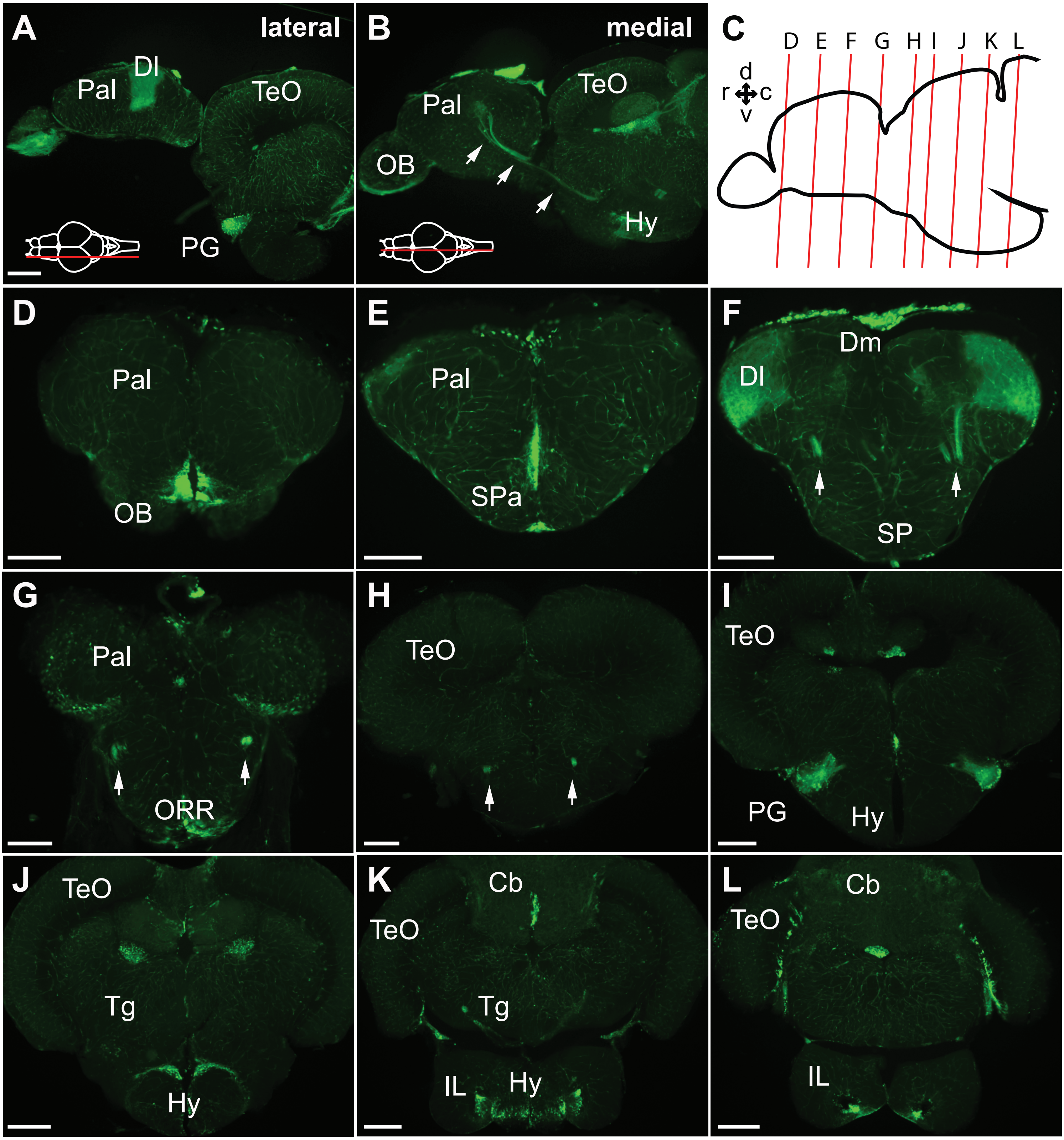

### Supplementary file 2

#### Supplementary file 2

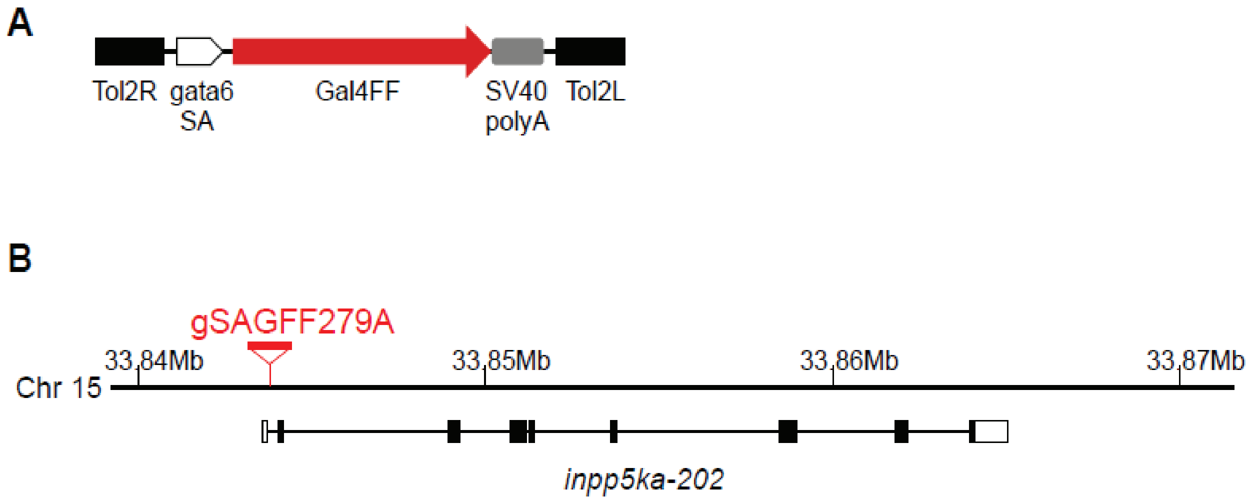

### Supplementary file 3

#### Supplementary file 3

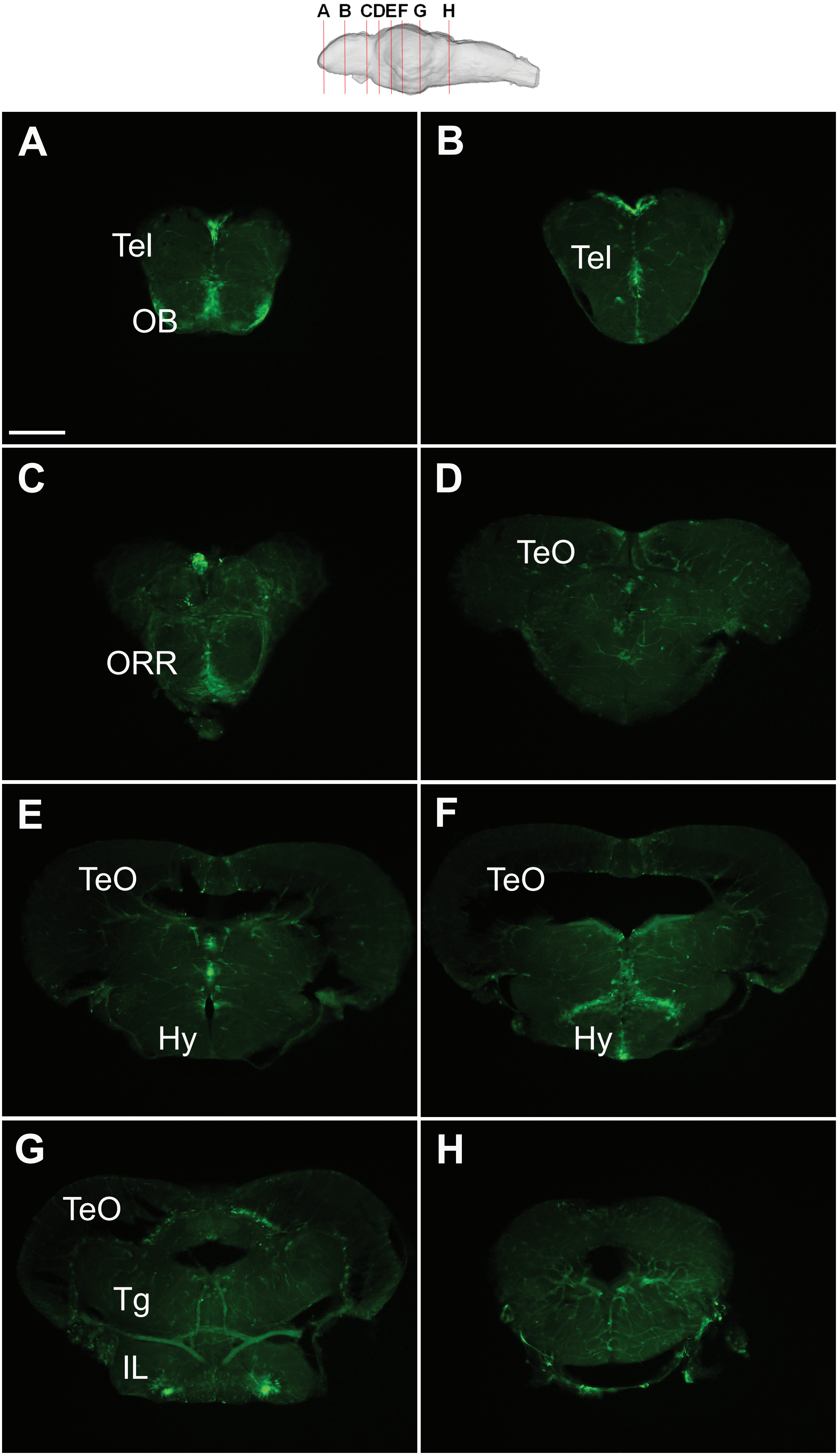

### Supplementary file 4

**Supplementary file 4**

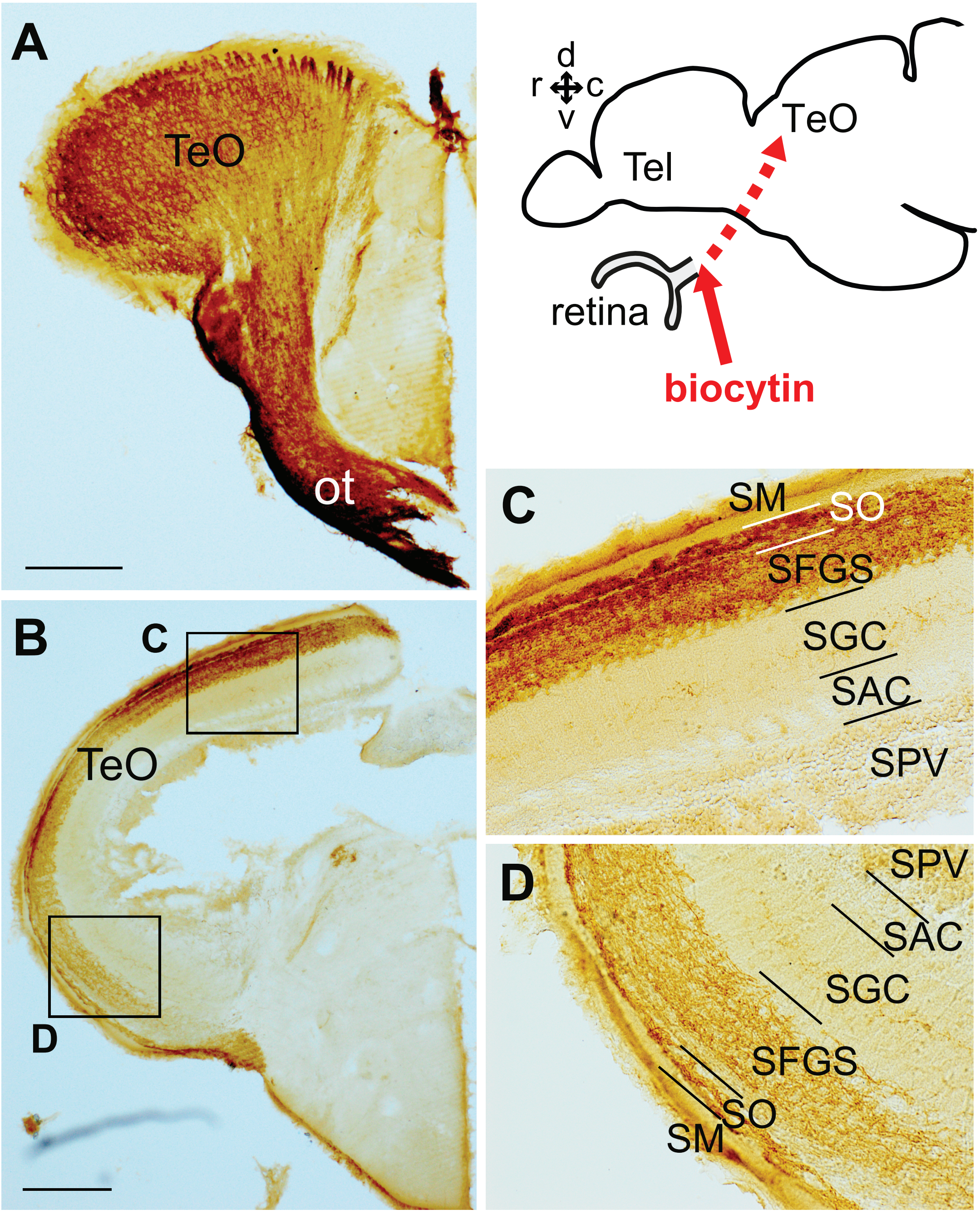

### Supplementary file 5

#### Supplementary file 5

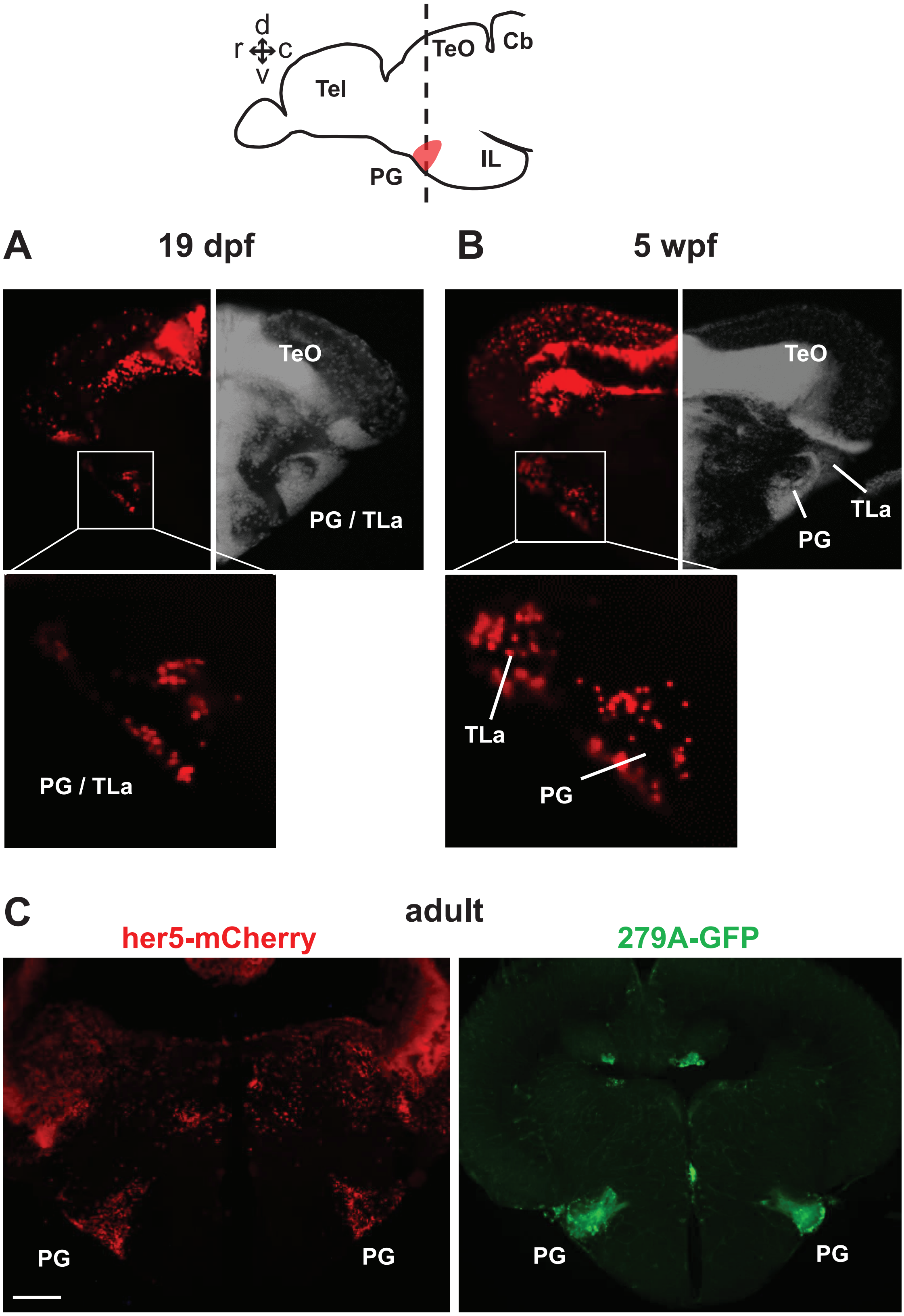

### Supplementary file 6

#### Supplementary file 6

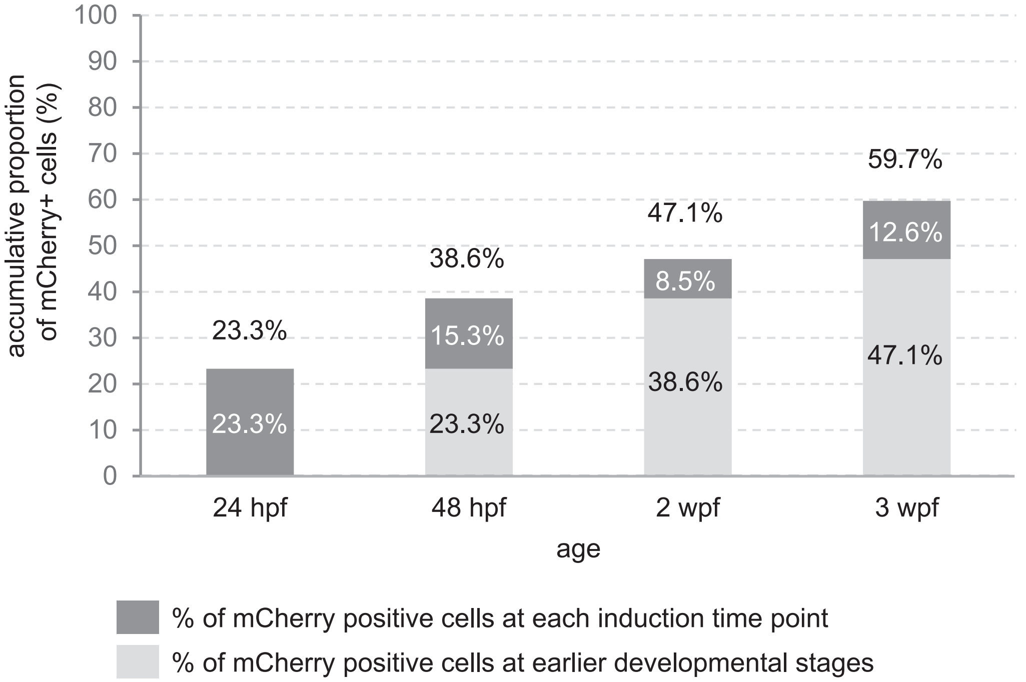

### Supplementary file 7

#### Supplementary file 7

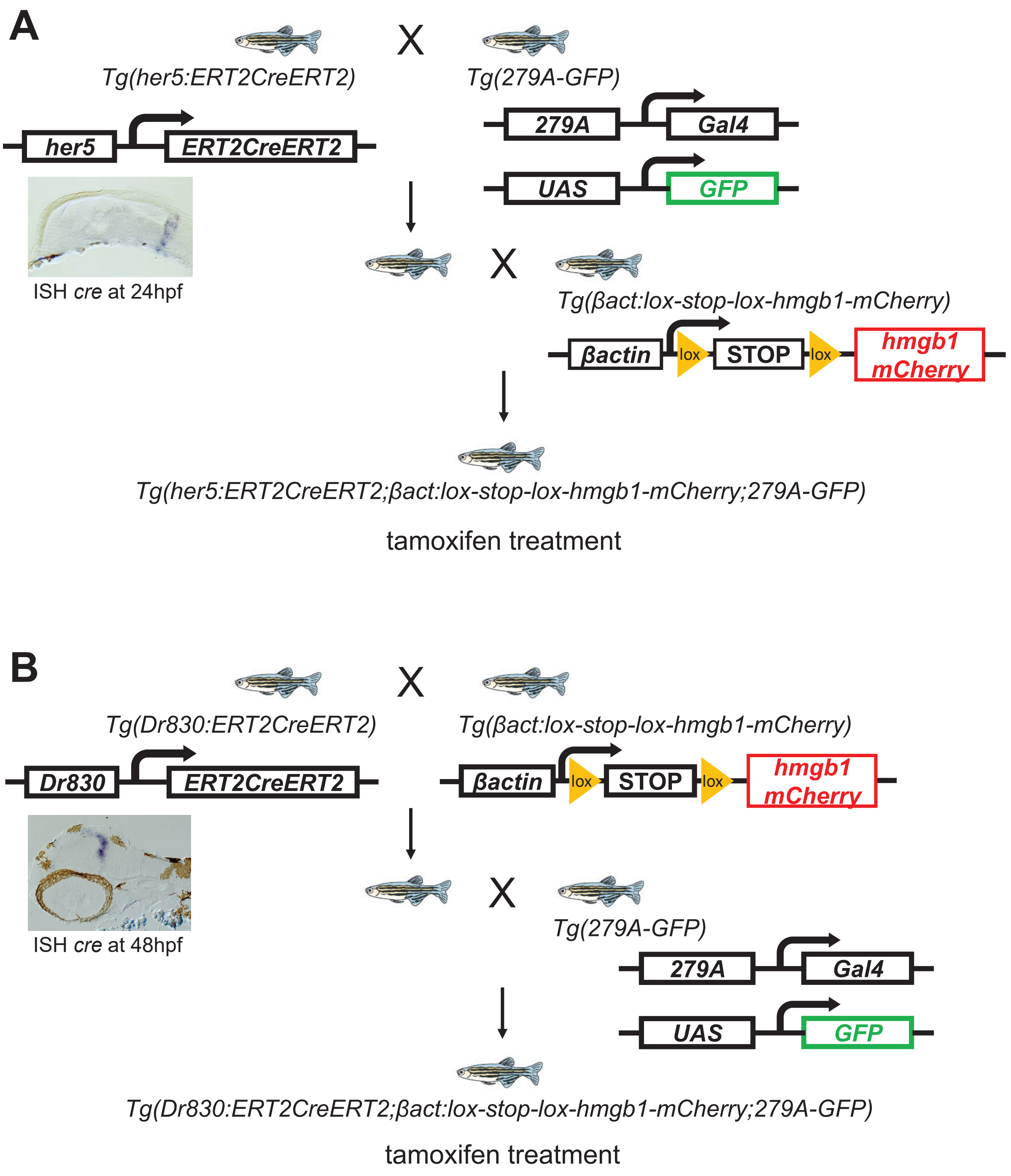

### Supplementary file 8

#### Supplementary file 8

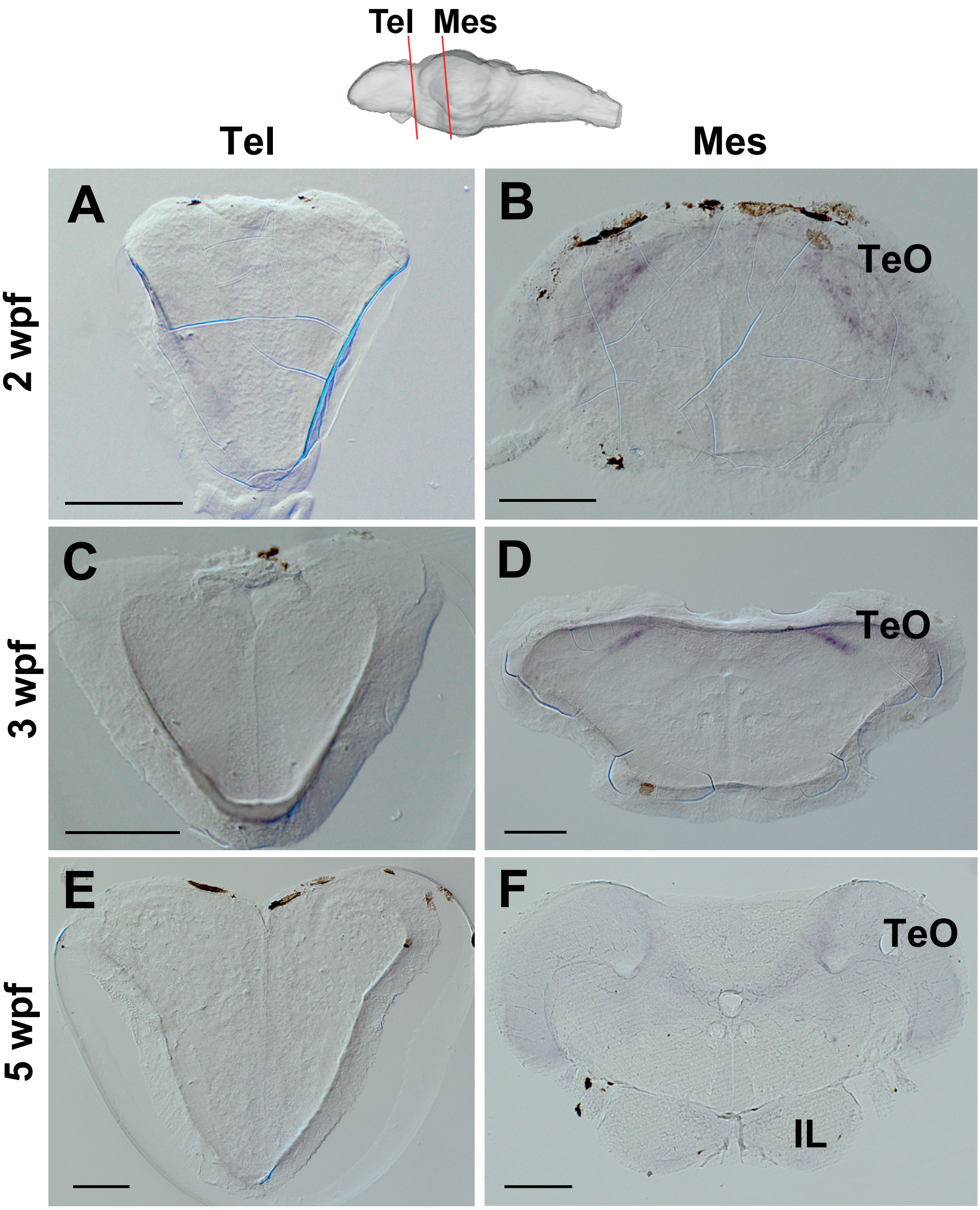
