## Supplementary file 9 for "Non-thalamic origin of zebrafish sensory relay nucleus: convergent evolution of visual pathways in amniotes and teleosts"

Supplementary file 9. Conditions of tamoxifen treatments

| age | | tamoxifen treatment | | | tamoxifen concentration | age of analysis | # fish analyzed |
|---|---|---|---|---|---|---|---|
| start | end | duration per day | number of days | total duration | | | |
| Tg(her5:ERT2CreERT2;βact:lox-stop-lox-hmgb1-mCherry;279A-GFP) | | | | | | | |
| 24 hpf | 30 hpf | 6 h | 1 | 6 h | 10 µg/mL | 3 mpf | 14 |
| Tg(Dr830:ERT2CreERT2;βact:lox-stop-lox-hmgb1-mCherry;279A-GFP) | | | | | | | |
| 24 hpf | 30 hpf | 6 h | 1 | 6 h | 10 µg/mL | 3 mpf | 3 |
| 24 hpf | 30 hpf | 6 h | 1 | 6 h | 10 µg/mL | 6 mpf | 6 |
| 30 hpf | 54 hpf | 24 h | 1 | 24 h | 5 µg/mL | 3 mpf | 6 |
| 48 hpf | 54 hpf | 6 h | 1 | 6 h | 10 µg/mL | 3 mpf | 6 |
| 3 dpf | 4 dpf | 24 h | 1 | 24 h | 5 µg/mL | 3 mpf | 6 |
| 7 dpf | 9 dpf | 4 h | 2 | 8 h | 5 µg/mL | 3 mpf | 6 |
| 2 wpf | 2 wpf | 2 h | 4 | 8 h | 2 µg/mL | 3 mpf | 6 |
| 3 wpf | 3 wpf | 2 h | 4 | 8 h | 2 µg/mL | 3 mpf | 6 |
| 4 wpf | 4 wpf | 2 h | 4 | 8 h | 2 µg/mL | 3 mpf | 6 |
| 5 wpf | 5 wpf | 2 h | 4 | 8 h | 2 µg/mL | 3 mpf | 6 |
| 6 wpf | 6 wpf | 4 h | 4 | 16 h | 2 µg/mL | 3 mpf | 6 |
| 8 wpf | 8 wpf | 4 h | 4 | 16 h | 2 µg/mL | 3 mpf | 6 |
